# Analysis of viral RNA-host protein interactomes enables rapid antiviral drug discovery

**DOI:** 10.1101/2021.04.25.441316

**Authors:** Shaojun Zhang, Wenze Huang, Lili Ren, Xiaohui Ju, Mingli Gong, Jian Rao, Lei Sun, Pan Li, Qiang Ding, Jianwei Wang, Qiangfeng Cliff Zhang

**Author notes:** Co-first author (Shaojun Zhang, Wenze Huang, Lili Ren, Xiaohui Ju, Mingli Gong, Jian Rao). Correspondence (Q.D.), (J.W.), (Q.C.Z.).

## Abstract

RNA viruses including SARS-CoV-2, Ebola virus (EBOV), and Zika virus (ZIKV) constitute a major threat to global public health and society. The interactions between viral genomes and host proteins are essential in the life cycle of RNA viruses and thus provide targets for drug development. However, viral RNA-host protein interactions have remained poorly characterized. Here we applied ChIRP-MS to profile the interactomes of human proteins and the RNA genomes of SARS-CoV-2, EBOV, and ZIKV in infected cells. Integrated interactome analyses revealed interaction patterns that reflect both common and virus-specific host responses, and enabled rapid drug screening to target the vulnerable host factors. We identified Enasidenib as a SARS-CoV-2 specific antiviral agent, and Trifluoperazine and Cepharanthine as broad spectrum antivirals against all three RNA viruses.

**One Sentence Summary:** Interactome analyses of host proteins and the SARS-CoV-2, EBOV, and ZIKV RNA genomes unveil viral biology and drug targets.

## Introduction

It is notable that three of the most dangerous infectious diseases of recent times are all caused by RNA viruses: SARS-CoV-2, ZIKV and EBOV. The SARS-CoV-2 coronavirus has caused the on-going Coronavirus Disease 2019 (COVID-19) pandemic, resulting in more than 20 million infections and 800 thousand deaths and global disruption of our society and economy(*1, 2*). Since its outbreak in 2012, ZIKV has infected millions of people(*3*), and has remained on the 2018 Blueprint list of priority diseases by the World Health Organization (WHO, February, 2018). EBOV has led to epidemics in Africa and has spread to Europe and North America; Ebola virus disease is severe with an average case fatality rate of ∼80% in past outbreaks(*4*).

RNA viruses typically have a relatively small genome and proteome(*5*), and therefore rely heavily on interactions with host factors to complete their life cycles(*6*). RNA binding proteins function on many aspects of cellular and viral processing, e.g., RNA translation, stabilization, modification and localization(*7, 8*), as well as anti-infection response against pathogens(*9*). Many studies have focused characterization of viral protein-host protein interactions(*10–15*). In contrast, the interactions between host proteins and viral RNA (vRNA) are much less well understood, despite the known importance of the viral RNA genome for multiple processes during infection, including viral genome translation and replication(*16*).

Recent years have seen an explosion of high-throughput methods that, in combination with diverse crosslinking reagents and data processing workflows, enable a global analysis of RNA-protein interactions (“the interactome”) in cells(*17, 18*). These approaches can substantially advance our understanding of the infection and pathology of RNA viruses and can inform diverse and effective therapeutic options, yet they have not yet been widely deployed for comparative analyses of cells infected with diverse RNA viruses or strains.

Here, we unveil the vRNA-host protein interactomes of SARS-CoV-2, EBOV, and ZIKV based on ChIRP-MS analysis of infected human host cells. Our integrated interactome analysis identified interaction patterns that reflect both common and virus-specific host responses. We applied these insights to inform a targeted antiviral drug screening workflow based on standard and/or trans-complementation viral infection assay systems that ultimately identified the FDA approved drug Enasidenib, which inhibits the SARS-CoV-2 specific interacting protein IDH2, as a potent SARS-CoV-2 specific antiviral agent. Further, we found that the heat shock protein inhibitor Cepharanthine and the cytoskeleton disruptor Trifluoperazine are broad spectrum antivirals against all three of these pathogenic RNA viruses.

## Results

### ChIRP-MS reveals the SARS-CoV-2 vRNA-host protein interactome

To define host proteins associated with genomic RNA of SARS-CoV-2, we used the RNA-directed proteomic discovery method ChIRP-MS (Comprehensive identification of RNA-binding proteins by mass spectrometry)(*19*) (Materials and Methods). Briefly, human hepatocarcinoma Huh7.5.1 cells were infected with SARS-CoV-2 and crosslinked with formaldehyde to preserve vRNA-protein complexes. Biotinylated oligonucleotides tiling SARS-CoV-2 vRNA with specificity were used to enrich vRNA-host protein complexes from cell lysates (Table S1) and the co-purified proteins were identified by mass spectrometry (Fig. 1A).

**Fig. 1.**
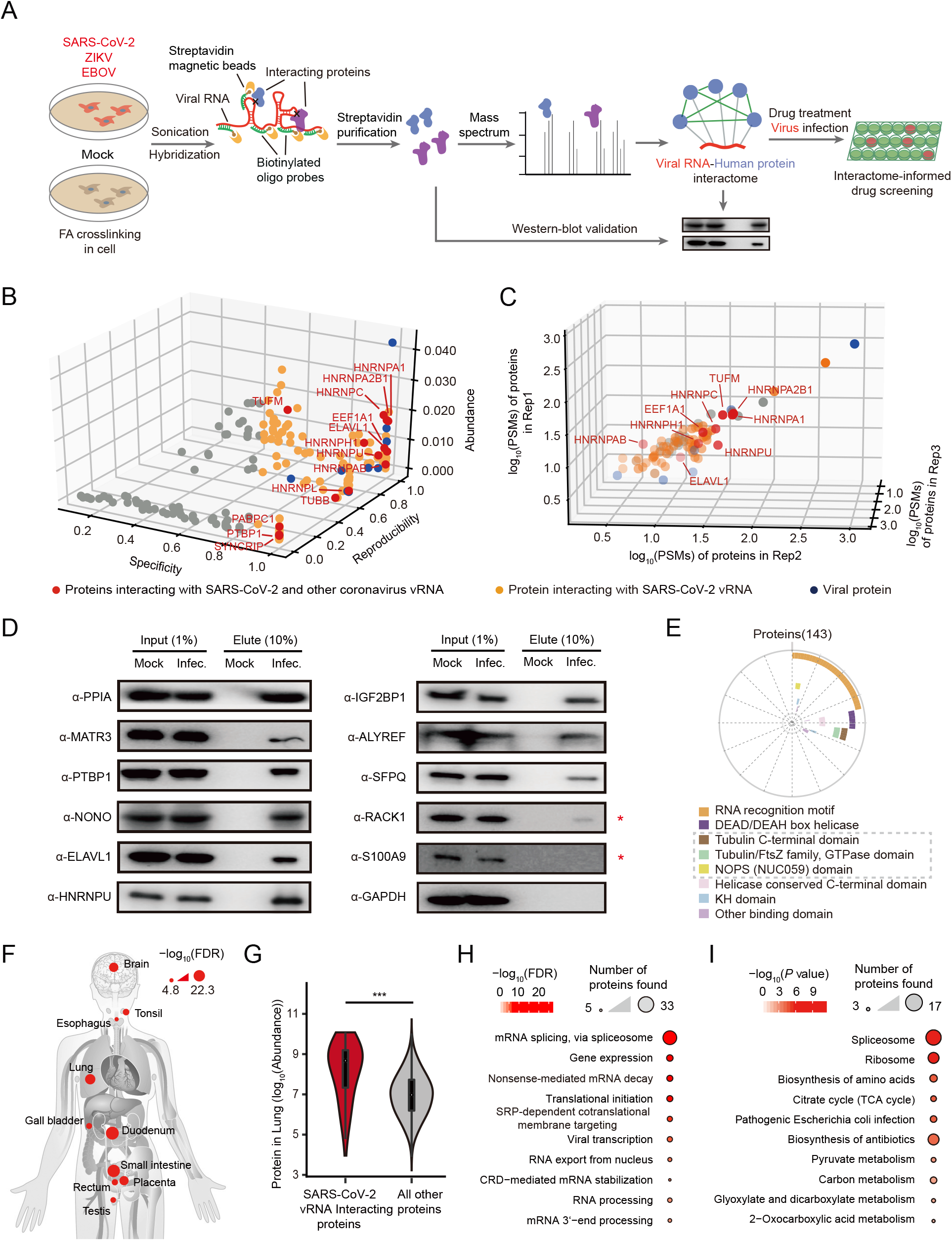
Identification of the human proteins interacting with the SARS-CoV-2 RNA genome in infected cells. **A**, Schematic for identifying host proteins that interact with viral RNAs using ChIRP-MS. FA, formaldehyde. ZIKV, zika virus. EBOV, Ebola virus. Human cells were infected with SARS-CoV-2, ZIKV or EBOV. Cells without infection (Mock) were used for control. Infected and mock cells were crosslinked using formaldehyde, and then sonicated to release RNA-protein complexes. For each virus, the viral RNA (vRNA)-human protein complexes were purified using biotinylated oligos specifically tiling the viral RNA genome, and the co-purified human protein were identified using mass spectrometry. **B**, Evaluation of the SARS-CoV-2 vRNA interacting proteins identified by ChIRP-MS. Peak intensities of peptides of a protein from the mass spectrum (Abundance), protein abundance over different replicates (Reproducibility), and uniqueness of an interacting protein across all replicated experiments (Specificity) were evaluated to define viral RNA interacting proteins (see Materials and Methods, ChIRP-MS data analysis). **C**, Correlation of log_10_ (PSMs) for proteins from three biological replicates. In **B** and **C**, interacting human proteins are indicated as red dots if also known to interact with other coronavirus genome RNA, and otherwise as orange dots; SARS-CoV-2 viral proteins co-purified with viral RNA are blue dots. **D**, Validation of the interactions between the SARS-CoV-2 vRNA and human proteins using ChIRP-Western blotting (ChIRP-WB). GAPDH protein, which is not known to bind SARS-CoV-2 vRNA, was used as a negative control for ChIRP-WB experiments. Mock, cells without infection. Infec., cells infected with SARS-CoV-2. *, indicates proteins that are not consistent between ChIRP-MS and ChIRP-WB. **E,** Distribution of enriched protein domains among the interacting human proteins of SARS-CoV-2 vRNA. Bold lines with different colors represent different protein domains. Protein domains not known as RNA binding domains are indicated using grey dashed boxes. **F,** Human tissues or organs with significantly elevated expression levels for SARS-CoV-2 vRNA interacting proteins (n=143). *P* value was calculated by Mann–Whitney U test and adjusted by FDR. The 10 tissues with the most significant *P* values are shown. **G,** SARS-CoV-2 vRNA interacting proteins are more abundant than other non-interacting proteins in lungs. *P* value was calculated by Mann–Whitney U test and adjusted by FDR. **H,** Gene Ontology enrichment analysis of the SARS-CoV-2 vRNA interacting proteins. The top 10 enriched terms are shown. **I,** Cellular pathway (KEGG) enrichment analysis of the SARS-CoV-2 vRNA interacting proteins. The top 10 enriched terms are shown.

ChIRP-MS analysis of SARS-CoV-2-infected Huh7.5.1 cells identified a total of 143 human proteins with a MiST score > 0.6 (Fig. 1B and Table S2). Note that the co-purification steps of the ChIRP-MS strategy depleted highly abundant host RNAs and enabled robust recovery of over 80% of the total viral RNA in infected cells (Fig. S1A). Correlation coefficients across three biological replicates were all above 0.9 (Fig. 1C). Our ChIRP-MS dataset underscored the extensive interactions between viral RNA and the SARS-CoV-2 nucleocapsid protein (NP), the viral M protein, the spike protein and several non-structural proteins (NS) (Table S2. Also see Fig. 1B), consistent with previous studies showing that NP directly binds and encases viral RNA during infection.

To validate the vRNA-interacting host proteins from the mass spectrometry based identification, we assessed the ChIRP samples using western blotting (ChIRP-WB) (Fig. 1A. Materials and Methods). We examined 11 candidate proteins using antibodies that were available in our lab (*e.g.*, HnRNPU and PPIA with a high MiST score 1, and RACK1 with a lower MiST score 0.68), and confirmed that 9 of them as SARS-CoV-2 vRNA interacting proteins in infected Huh7.5.1 cells (Fig. 1D). Among them, ELVAL1, hnRNPU, and PTBP1 were previously revealed to bind other coronavirus vRNAs that function in viral replication(*20, 21*). PPIA (Cyclophilin A) has been widely implicated in viral infection, and drugs targeting PPIA showed broad-spectrum antiviral activity(*22*). MATR3 may associate with SFPQ/NONO heterodimers to modulate the STING pathway(*23*). IGF2BP1 is a well-characterized RNA binding protein involved in RNA stability and viral RNA translation(*24*). The THO complex subunit protein (ALYREF) is a component of the TREX complex that is involved in RNA processing and transport(*25*). Offering further support that our approach yielded biologically relevant candidates rather than processing-related artefacts, we noted that the vRNA-interacting candidate host proteins were enriched for canonical RNA-binding domains (*e.g.*, RNA recognition motif, RNA secondary structure related DEAD/DEAH box helicase domain, and RNA-binding KH domain) (Fig. 1E, Fig. S1B).

We comapared our 143 SARS-CoV-2 vRNA binding proteins with host factors previously reported to bind coronaviruses, and found that 14 of the host factors in our ChIRP-MS interactome are known to bind the viral RNA of other coronavirus and to impact viral replication(*20, 21*) (Fig. 1B). In particular, we comapared our dataset with a recently reported host interactome for SARS-CoV-2 based on the “RNA antisense purification and mass spectrometry” (RAP-MS) method(*26*), and found 17 host factors common to both interactome datasets (Fig. S1C). In addition, of the 143 SARS-CoV-2 host RNA binding proteins that we identified, 69 (61%, Table S2) were previously reported as responsive to SARS-CoV-2 infection in a proteomics study(*27*). Collectively, these findings suggest that ChIRP-MS can reliably identify vRNA-interacting host factors to thus to catalog a high-confidence SARS-CoV-2 vRNA interactome.

SARS-CoV-2 infects multiple organs, leading to diverse symptoms such as fever, shortness of breath, severe acute pneumonia, haempotysis, diarrhea, and neurological disorders(*28*). Indeed, we found that many of the SARS-CoV-2 vRNA-binding proteins were more highly expressed in organs known to host SARS-CoV-2 infection, such as lung(*29*), intestines(*30*) and brain(*31*) (Fig. 1F-G). We also performed Gene Ontology (GO) and Kyoto Encyclopedia of Genes and Genomes (KEGG) analyses and found that the annotations of the interactome host factors were enriched for GO terms related to splicing, translation, and viral transcription (Fig. 1H), as well as KEGG pathways related to the spliceosome, ribosome and RNA metabolism (Fig. 1I). The aforementioned proteomics survey also reported that host proteins responsive to SARS-CoV-2 infection are similarly enriched for annotated functions relating to translation, RNA metabolism, and RNA transport related cellular processes(*27*).

### High-confidence vRNA-human protein interactomes for EBOV and ZIKV

Seeking to delineate which host proteins are common to different RNA viruses vs. virus-specific factors, we used ChIRP-MS to uncover the vRNA-host protein interactomes for the ZIKV MR766 strain and a recombinant EBOV Δ VP30-GFP virus with the VP30 coding sequence of the Ebola virus (strain Zaire Mayinga) replaced by the GFP sequence(*32*) (Materials and Methods). As above, we employed biotinylated oligonucleotides tiling each viral RNA with specificity to enrich for viral RNA-host protein complexes from lysates of infected cells (Table S1), and identified the co-purified proteins by mass spectrometry (Fig. 1A).

These ChIRP-MS analyses identified a total 172 viral RNA binding proteins in MR766-infected Huh7 cells (MiST score > 0.7) and 223 viral RNA binding proteins in EBOVΔVP30-GFP infected Huh7.5.1-VP30 cells (MiST score > 0.6, Fig. 2A-B and Table S2). In each analysis, the co-purification steps of the ChIRP-MS strategy allowed for the depletion of highly abundant host RNAs, and, as with SARS-CoV-2, enabled robust recovery of more than 80% of the total viral RNA in infected cells (Fig. S2A). All data showed high correlations between the experimental replicates (*r* > 0.7 and 0.9 respectively for MR766 and EBOV, Fig. 2B-C).

**Fig. 2.**
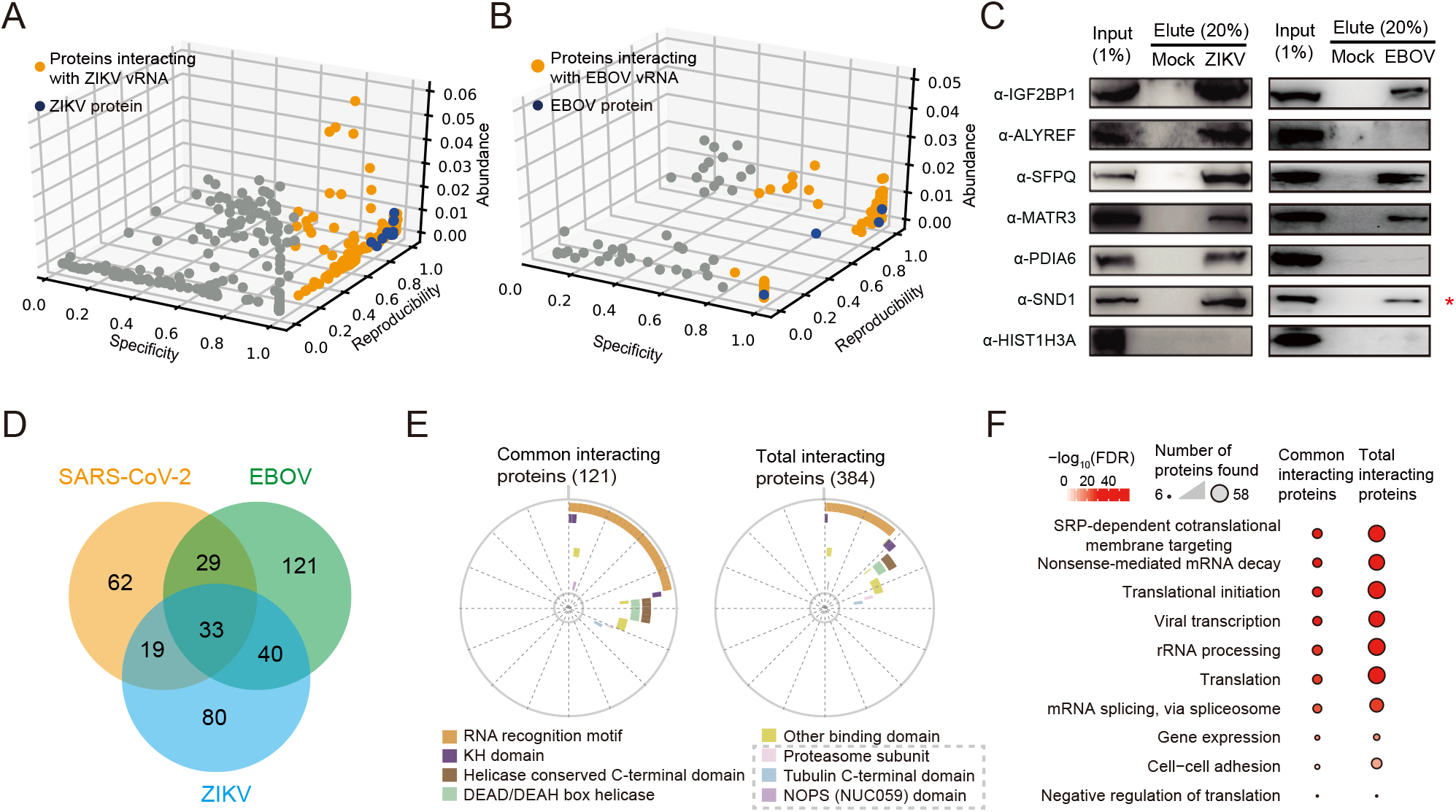
ChIRP-MS identified the human proteins interacting with the ZIKV and EBOV RNA genomes in infected cells. **A, B**, Evaluation of the ZIKV **(a)** or EBOV **(b)** vRNA interacting proteins identified by ChIRP-MS. Definitions of “specificity”, “abundance” and “reproducibility” are the same as Fig. 1B (see Materials and Methods, ChIRP-MS data analysis). Host proteins interacting with the ZIKV or EBOV vRNA are indicated using orange dots. ZIKV or EBOV proteins co-purified with viral RNA are indicated as blue dots. **C,** Validation of interactions between the indicated proteins with ZIKV or EBOV vRNA using ChIRP-WB. HIST1H3A was used as a negative control. Mock, cells without infection. ZIKV or EBOV, cells infected with indicated virus. *, indicates proteins that are not consistent between ChIRP-MS and ChIRP-WB. **D,** Comparison of common interacting proteins between the viral RNA interactomes for SARS-CoV-2, EBOV, and ZIKV. **E,** Distribution of enriched protein domains among the common (left) and total (right) interacting human proteins of SARS-CoV-2, ZIKV, and EBOV vRNA. Bold lines with different colors represent different protein domains. Protein domains not known as RNA binding domains are indicated using grey dashed boxes. **F,** Gene Ontology enrichment analysis of the common (left) and total (right) interacting human proteins of SARS-CoV-2, ZIKV, and EBOV vRNA. The top 10 enriched terms are shown.

We again used ChIRP-WB to verify six of the host RNA binding proteins uncovered by ChIRP-MS for both ZIKV and EBOV (IGF2BP1, ALYREF, SFPQ, MATR3, PDIA6 and SND1) and validated their interactions with viral RNAs in ZIKV-infected Huh7 cells and in EBOV infected Huh7.5.1 cells (Fig. 2C). With only one exception (interaction between SND1 and EBOV vRNA), all validation results were consistent with our ChIRP-MS results. We then examined 12 host proteins that were shown previously to interact with various ZIKV strains, 8 of these 12 host proteins passed our scoring cutoff, suggesting a very high sensitivity for our ChIRP-MS experiments (Table S2). In addition, comparisons against previously reported ChIRP-MS based studies of host proteins that interact with another ZIKV strain(*17*) also revealed high level of overlap (Fig. S2D). Similarly, we also examined proteins that previously reported to interact with EBOV vRNA (DHX9, hnRNPR, hnRNPL, SYNCRIP, IGF2BP1)(*33*), and confirmed that all these interactions are covered in our EBOV ChIRP-MS result (Table S2). Overall, these analysis and observations suggest that our ChIRP-MS reliably captured host proteins that interact with the ZIKV and EBOV vRNAs in infected cells.

### An integrated analysis of the three RNA virus host factor interactomes identifies common and virus-specific host cellular factors and responses

Our interactome datasets highlight that the three RNA viruses interact with many common host proteins, but also showcase their virus-specific host interaction partners. In total, 33 human proteins interact with viral RNAs of all three viruses, and 88 proteins interact with viral RNAs of two of the three viruses, results suggesting the potential involvement of common host factors in conserved host reponses to infection by RNA viruses (Fig. 2D). Note that compared to the total set of all vRNA-interacting proteins, these common interacting proteins exhibit stronger enrichment for canonical RNA-binding domains (Fig. 2E), and consistently showed enrichment of cellular functions such as splicing, translation, and metabolism related processings (Fig. 2F).

To further characterize conserved host responses to RNA virus infection, we examined the common interacting protein complexes (and protein associations of the same biological pathways) shared by the three RNA viruses. We searched for enriched protein complexes and defined a complex as common one if it was present in at least two of the three virus interactomes (Fig. 3. Table S3. Materials and Methods). The analysis identified many common protein complexes, including the ribosome (and related translation regulators such as eIF4A), the spliceosome (and also other RNA processing proteins such as YBX1 and RBMX), the IGF2BP1-associated complex, the large drosha and DGCR8 complexes, the TNF-alpha/NF- κ B signaling complex, the microtubules and the cytoskeleton, the proteasome, and many stress-granule-related proteins.

**Fig. 3.**
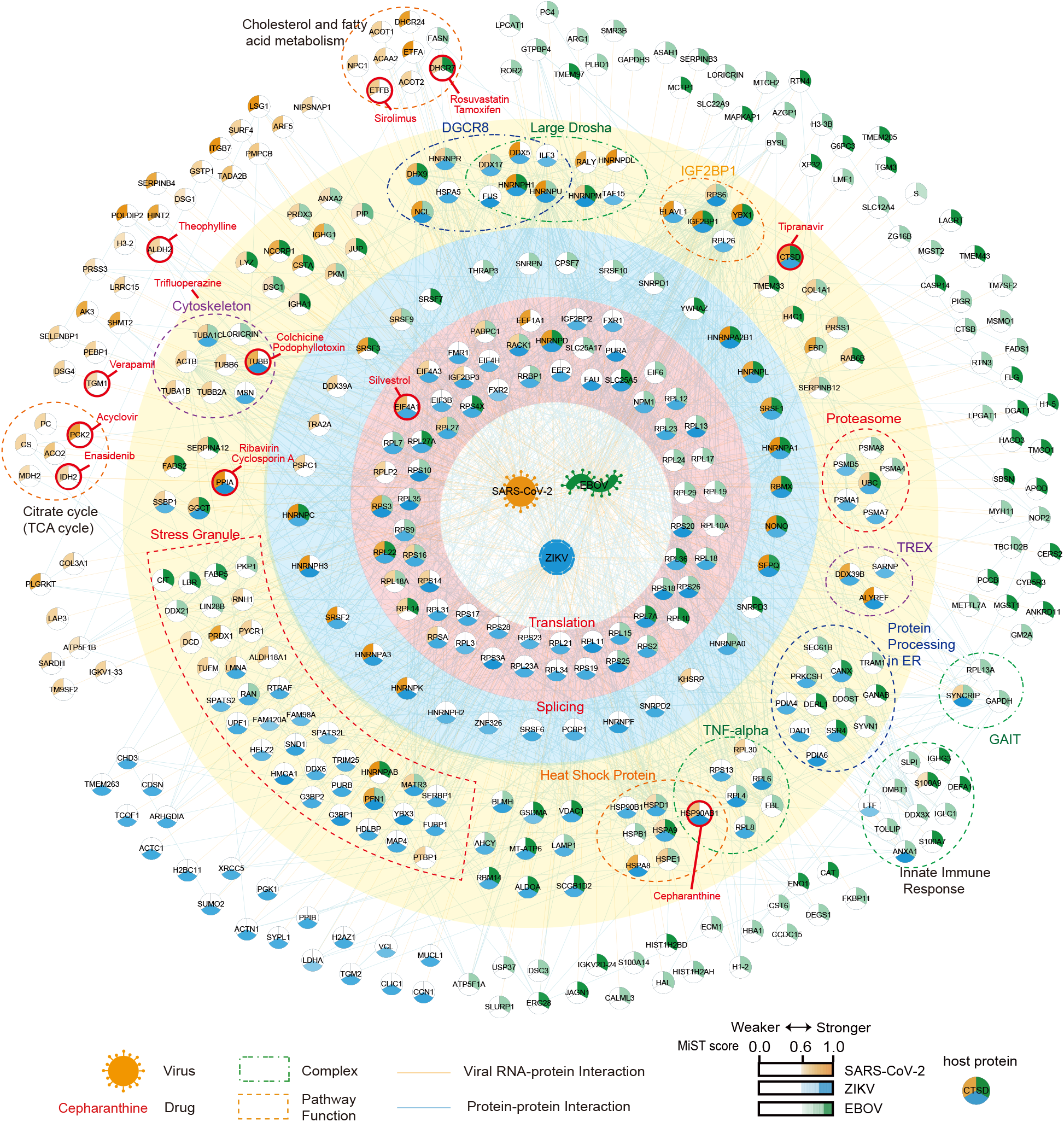
A compendium of viral RNA-human protein interactomes for SARS-CoV-2, EBOV, and ZIKV. Central nodes represent viral RNAs. Circular nodes represent host proteins. Edges indicate RNA-protein interactions (RPI, orange) and protein-protein interactions (PPI, blue). Protein complexes based on the CORUM database and proteins associated with the same biological processes according to functional annotation are enclosed with dashed lines. Nodes are colored according to the MiST scores of proteins interacting with SARS-CoV-2 (orange), ZIKV (blue), and EBOV (green) vRNA. Drugs targeting or predicted to target the indicated proteins (Table S5, Materials and Methods, Interactome-informed drug discovery) are highlighted with red circles.

Many of these common complexes are known to function in the regulation of host cell and/or viral RNA. For example, the IGF2BP1-associated complex regulates the stability and translation of various cellular RNA molecules including c-*Myc*(*34*) and also enhances IRES-mediated translation initiation of HCV RNA(*24*). The large drosha and DGCR8 complexes, which are involved in miRNA biogenesis, are essential for degrading exogenous double-stranded RNA, including viral RNAs(*35*). The cytoskeleton is often rearranged by viral infection and is involved in the formation of virus-induced membranous replication factories and that chemicals targeting the cytoskeleton are known to inhibit the replication of RNA virus(*36*). Of particular interests, host factors that function as stress-granule-related proteins(*37*), were repeatedly observed in the interactomes of all three viruses (Fig. 3), supporting the idea that stress granules as a general host cell response to viral infection.

Of equally paramount interest, we also identified many virus-specific protein complexes and pathways (Fig. 3, Fig. 3A-F, Table S3). For example, the SARS-CoV-2 vRNA interactome was specifically enriched for two metabolism pathways: the tricarboxylic acid (TCA)-cycle and the 2-Oxocarboxylic acid metabolism (*i.e.*, lipid metabolsim). Indeed, previous proteomic investigation also observed that these pathways are disrupted by SARS-CoV-2 infection(*27*). We also discovered that ZIKV interacts with components of the TREX transcription/export complex (comprising THOCs, ALYREF, DDX39B and SARNP, Fig. 3). This protein complex is important for RNA transport and its disruption has been implicated in microcephaly(*38, 39*), a disease associated with ZIKV infection(*40*). The SARS-CoV-2 host factor interactome includes only two known THOC complex subunits, while the EBOV includes none. We also found that EBOV specifically interacts with a set of immune and inflammatory responses related proteins(*41–43*), including S100A9, ANXA1, and DDX3X, and especially the “IFN-γ-activated inhibitor of translation” (GAIT) complex (Fig. 3), which is known to repress the translation of inflammatory related mRNAs(*44*). At minimum, these findings warrant further investigation for a functional connection to the organ failure and death that arise from the dysregulated host inflammatory responses during EBOV infection(*45*).

### Interactome-informed drug repurposing discovered potent SARS-CoV-2 antiviral compounds

The selective inhibition of host vRNA-binding proteins can attenuate infection and inform the development of innovative therapies(*12*). Accordingly, existing drugs that are known to target various host factor proteins can be exploited as potential treatments strategies during epidemics and pandemics. Given the urgent need for COVID-19 therapies(*46*), we analyzed open-source chemical databases to uncover compounds that target host factors within the SARS-CoV-2 vRNA interactome. Specifically, we investigated the IUPHAR/BPS Guide to Pharmacology, Drugbank, Drugcentral, and ChEMBL databases, and identified a total of 5309 compounds (Table S4) that are known or predicted to interact with 56 SARS-CoV-2 vRNA-binding proteins. We prioritized approved drugs and clinical phase agents, as well as those available at the Center of Pharmaceutical Technology of Tsinghua University. Ultimately, we tested the antiviral activities of 21 drugs (Fig. 4A, Table S5), including 20 FDA-approved drugs and 1 agent (Silvestrol) being tested in preclinical trials.

**Fig. 4.**
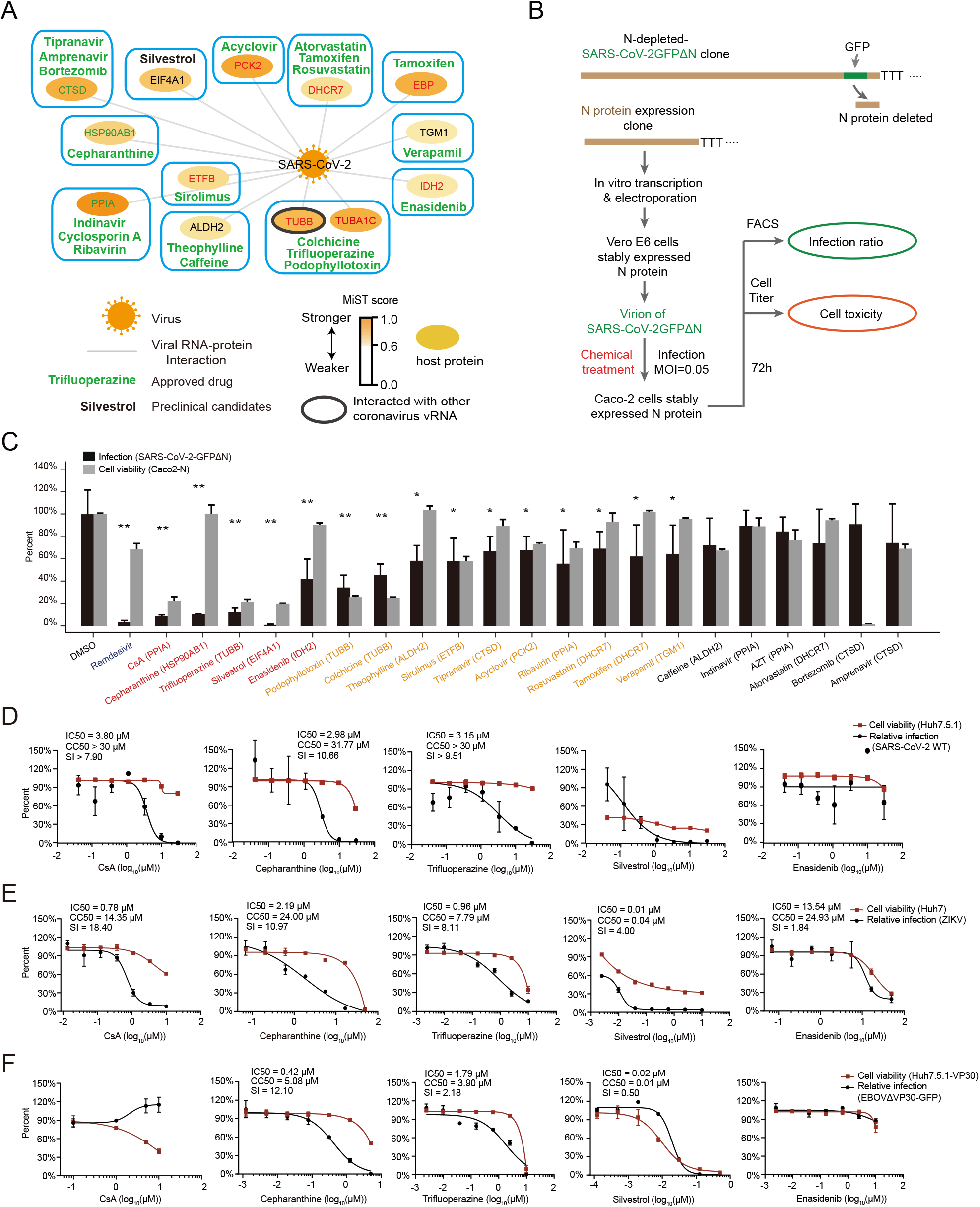
Validation of antiviral activities of the FDA-approved and clinical-trial drugs targeting viral RNA interacting proteins. **A,** FDA-approved (green) and clinical-trial (black) compounds modulating the SARS-CoV-2 vRNA interacting proteins are presented. Nodes are colored by the MiST scores of SARS-CoV-2 vRNA interacting proteins, and protein names are colored by the up- (red) or down-regulation upon SARS-CoV-2 infection(*27*). **B,** Antiviral activity test for the selected drugs using a SARS-CoV-2-GFPΔN trans-complementation system. For this validation system, the SARS-CoV-2 genome sequence segment encoding nucleocapsid (N) was replaced by GFP, thus the resulted SARS-CoV-2-GFPΔN only amplifies in N protein-expressing Caco-2 (Caco-2-N) Δ cells. The infection ratio was then evaluated by quantifying the GFP positive cells. For initial testing of antiviral activities, drugs were added at the same time as the infection (MOI=0.05), at a concentration of 10μM. GFP ratios (indicating the μ infection ratio of SARS-CoV-2) were calculated based on FACS data after 72 h. Remdesivir was used as an antiviral positive control. **C**, SARS-CoV-2-GFPΔN infection ratios in cells treated with the indicated drugs at a Δ concentration of 10μ Caco2-N cells were infected with SARS-CoV-2-GFPΔN virus for 72 h with drug treatment at day 0. Black bar, infection ratios of the SARS-CoV-2-GFPΔN. Grey bar, cell viabilities after treatment with the indicated compounds at 10 M. Data were normalized to DMSO treatment. Remdesivir as a μ positive control (2μ). Drugs selected for further validation were highlighted in red. Drugs with modest antiviral effects were highlighted in yellow. Drugs with no significant antiviral activities at a concentration of 10μ means ± SD. n= 4 biologically independent samples. **<0.01, *<0.05, Student’s*t*-test. **D-F,** The antiviral activities of Silvestrol, CsA, Cepharanthine, Trifluoperazine, and Enasidenib against SARS-CoV-2 **(D)**, ZIKV **(E)**, and EBOV Δ VP30-GFP **(F)** infections at different drug concentrations. Red line, cell viability; Black line, infection ratio relative to DMSO treatment. Data are means ± SD. n=3 biologically independent samples. IC50, CC50, and SI values are indicated.

To efficiently assess the potential antiviral effects on SARS-CoV-2 infection for these drugs, we used a trans-complementation system based on human colorectal adenocarcinoma epithelial Caco2 cells that supports infection and replication of SARS-CoV-2 (Fig. 4B). Briefly, this system uses a SARS-CoV-2-GFP-2 genome wherein the N protein-coding sequence is replaced with GFP; this replacement both disrupts the packaging capacity of the virus and supports visualization. SARS-CoV-2-GFP-2 vRNA was generated by *in vitro* transcription and transfected into green monkey epithelial Vero cells that stably express the SARS-CoV-2 N protein. These steps yielded packaged SARS-CoV-2-GFP-2 virions, which we used to infect Caco2 cells that also stably express the SARS-CoV-2 N protein (Caco2-N). We used GFP expression as a surrogate to monitor viral infection and replication in human cells.

We treated Caco2-N cells with each drug at the same time as the viral challenge. Remdesivir, which targets the viral RdRp and is well known for its antiviral activity against SARS-CoV-2, was used as a positive control, and vehicle (DMSO) was used as a negative control. At 72 h post infection we quantified infected, GFP-positive cells by FACS. Remdesivir (2µM) was found to strongly inhibit SARS-CoV-2-GFP-2 infection, with an infection about 3.7% relative to the negative control DMSO-treated cells (Fig. 4C). Thus, our trans-complementation system reflects SARS-CoV-2 replication in cells and is suitable for initial drug-testing.

Our initial screen using 10µM of each selected compound uncovered activity against SARS-CoV-2 for 15 of the 21 tested candidate drugs (Fig. 4C). Five exerted strong antiviral effects (Fig. 4C): 1) Silvestrol, which targets the translation initiation factor eIF4A(*47*) (preclinical on Chronic lymphocytic leukemia; 98.8% inhibitory efficiency); 2), Cyclosporin A (CsA), which is an immunosuppression drug and targets protein PPIA(*48*) (FDA-approved for immunosuppressant; 91.2% inhibitory efficiency); 3), Trifluoperazine (FDA-approved for antipsychotic and an antiemetic; 87.5% inhibitory efficiency) disrupts cytoskeleton organization(*49*) and also inhibits binding between HSP90 with its chaperons(*50*), and 4) Cepharanthine (Approved for alopecia, 89.6% inhibitory efficiency), which is an inhibitor of TNF-alpha/NF-κB, and also an inhibitor of the heat shock proteins (HSP90s and HSPA8)(*51, 52*) and 5) Enasidenib, which targets the mitochondrial energy related enzyme IDH2(*53*) (FDA-approved for refractory acute myeloid leukemia; 58% inhibitory efficiency).

Despite the potential for inhibition of SARS-CoV-2 infection at the initial 10μM concentrations, these five candidate drugs also showed varying levels of cytotoxicity (Fig. S4A). We further used the trans-complementation assays to determine the antiviral potency and cytotoxicity at different concentrations for the aforementioned five candidates (Fig. 4D). For testing the drug effects, we infected Caco-2 cells with SARS-CoV-2 virus (MOI 0.05) and treated cells with the compounds with different concentrations. Remdesivir, as positive control, reduced the infection of SARS-CoV-2 M (Fig. S4B). Silvestrol was previously reported as a μ “broad-spectrum” antiviral agent that showed inhibition activity against the coronavirus HCoV-229E(*54*). Although Silvestrol showed a dose-dependent effect on SARS-CoV-2 infection (IC50=0.32µM), we found that the cell viabilities dropped dramatically at higher drug concentrations, and detected no inhibitory effects on infection of SARS-CoV-2 virus infection below a 100nM of Silvestrol concentration. CsA can inhibit infection of SARS-CoV-2 virus with an IC50 value of 3.8µM, and a CC50 value > 30µM in Caco-2 cells (SI > 7.9). Cepharanthine can inhibit SARS-CoV-2 infection with an IC50 value of 2.98µM, and a CC50 value of 31.77µM (SI=10.66). Trifluoperazine also showed strong antiviral activities against SARS-CoV-2, with IC50 value of 3.15µM, and CC50 value > 30µM (SI > 9.51). Enasidenib, targeting the SARS-CoV-2 vRNA specific host factor IDH2, showed no antiviral effects. Thus, among the tested candidate drugs, Cepharanthine (approved in Japan) and Trifluoperazine (FDA-approved) are the most attractive for potential use for against SARS-CoV-2 infection.

### Identification of broad antiviral inhibitors against multiple RNA viruses by targeting common host factors

Our viral RNA interactome data indicated that both ZIKV and EBOV vRNAs interact with the translational machinery, heat shock proteins (*e.g.*, HSP90s, HSPA9, HSPA8 HSPD1, etc.), as well as cytoskeleton-related proteins (*e.g.*, TUBA1C, TUBB). This overlap amongst host factors for these RNA viruses motivated us to examine the antiviral effects of drugs targeting these host proteins, specifically using infection assays for ZIKV in Huh7 cells (MR766, MOI 0.5) and for EBOV using assays in in Huh7.5.1-VP30 cells based on an EBOVΔVP30-GFP variant(*32*) (Fig. 4E-F).

The translation initiation inhibitor Silvestrol inhibited ZIKV infection with an IC50 value of 0.01µM and a CC50 value of 0.04 µM (SI=4). CsA inhibited ZIKV infection with an IC50 value of 0.78µM, and a CC50 value of 14.35µM (SI=18.40). And the cytoskeleton-disrupter Trifluoperazine inhibited ZIKV infection with an IC50 value of 0.96µM, and a CC50 value of 7.79µM (SI=8.11). The most potent anti-ZIKV effects we observed in these assays was from Cepharanthine that targets heat shock proteins, which inhibited ZIKV infection with an IC50 value of 2.19µM, and a CC50 value of 24µM (SI=10.97) (Fig. 4E). And Enasidenib targeting IDH2 showed low inhibitory effects on ZIKV infection (SI = 1.84).

As a negative-strand RNA virus, the infection and replication processes of EBOV are very distinct from those of positive-strand RNA virus such as SARS-CoV-2 and ZIKV(*4*). Consistently, our integrated viral RNA interactome analysis for the three RNA viruses indicated that SARS-CoV-2 and ZIKV share more overlapped host factors than that of any with EBOV, specifically, PPIA interacts with vRNAs of SARS-CoV-2 and ZIKV, while not with vRNA of EBOV. Indeed, the PPIA-inhibitor CsA showed antiviral activities against SARS-CoV-2 and ZIKV but not EBOVΔVP30-GFP viral infection (MOI 0.1), even at a high concentration (10 µM, Fig. 4D-F). This result is also consistent with a previous study reporting that another PPIA inhibitor (Alisporivir) has limited antiviral activity against EBOV(*55*). Again, Silvestrol showed obvious cell toxicity although it exerted dose-dependent inhibition to EBOVΔVP30-GFP infection (IC50 = 0.02 µM), and Trifluoperazine (IC50 = 1.79 µM, SI = 2.18) and Cepharanthine (IC50 = 0.42µM, SI = 12.10) showed antiviral activities to EBOVΔVP30-GFP virus infection. The IDH2 inhibitor Enasidenib also showed no effects on EBOVΔVP30-GFP virus infection. Note that in our study here, we used SARS-CoV-2 for initial test of the antivirals, which could potentially miss EBOV specific antivirals, given the distinct life cycle of positive-strand and negative-strand RNA viruses. It is worthy in the future to perform an EBOV host factor guided-large scale screening for discovery of more effective antivirals against EBOV.

Collectively, our results support that the FDA-approved cytoskeleton disrupter drug Trifluoperazine and the heat shock proteins inhibitor Cepharanthine have broad-spectrum antiviral activity against the three major RNA viruses.

## Conclusions

Our ChIRP-MS profiling of the vRNA-host protein interactomes of human cells infected with the causal pathogenic viruses for COVID-19, Zika and Ebola virus diseases identified interaction patterns that reflect both common and virus-specific host responses. These interactome datasets can be viewed as a rich resource to help illuminate the infection biology for these three viral pathogens and potentially others as well. The RNA-centric view of ChIRP-MS was particularly informative in revealing molecular virology of all stages of the life cycle of RNA viruses in infected cells(*17*). Methods to analyze RNA-protein interactome datasets are not yet standardized, and we found that integrated comparisons offered a straightforward way to explore and visualize common and virus-specific interactions, which we arranged to highlight the functions of the various host proteins. Indeed, the analysis enabled the prediction and selection of small molecule inhibitors as either specific or likely broad-spectrum antiviral agents. And finally, we were able to combine the interactome-informed predictions and an innovative trans-complementation infection system to power a pipeline for rapid antiviral drug discovery.

Ultimately, these efforts demonstrated that the FDA-approved IDH2 inhibitor Enasidenib is a potent SARS-CoV-2 specific antiviral agent and that agents targeting the heat shock proteins and cytoskeleton components, Cepharanthine and Trifluoperazine, can exert broad-spectrum effects against all three RNA viruses we examined in our study. Having identified the activities of Enasidenib, Cepharanthine, and Trifluoperazine, next steps for their re-purposing and deployment as possible clinical antiviral agents would be *in vivo* infection model studies and pre-clinical safety evaluations. Moreover, deeper exploration of the antiviral modes of action for these drugs should reveal biological insights about how these viruses infect cells, how host cells respond, and which (if any) countermeasures the viruses may deploy to overcome host defense pathways. The strong current focus of the biomedical research community on the biology of RNA virus infection, especially the ongoing COVID-19 pandemic(*56*), has exposed how little we know about the infection and amplification mechanisms that these viruses employ. Our work illustrates how profiling datasets obtained with ChIRP-MS can be integrated to drive both basic discovery and therapeutic applications.

## Acknowledgments

We thank members of the Zhang lab for discussion. We thank professor Jianbin Wang for the help of the project. We thank the Tsinghua University Branch of China National Center for Protein Sciences (Beijing) for the computational facility support.

## Author contributions

Q.C.Z. conceived the project. Q.C.Z., J.W. and Q.D. supervised the project. L.R. prepared the SARS-CoV-2 virus assisted by J.R.. S.J. performed the ChIRP-MS and validation experiments assisted by L.S.. X.J. developed the SARS-CoV-2 N trans-complementation system and screened the drugs. L.G. screened the drugs on EBOV. W.H. analyzed the ChIRP-MS data with the assisted by P.L.. Q.C.Z. and S.Z. wrote the manuscript with inputs from all authors.

## Supplementary Materials

### Materials and Methods

#### Data reporting

No statistical methods were used to predetermine sample size. The experiments were not randomized and the investigators were not blinded to allocation during experiments and outcome assessment.

#### Cell line and antibodies

Cell lines and antibodies used in this study are listed in (Table S6). All cell lines were determined to be free of mycoplasma based on PCR and nuclear staining. All cell lines mentioned in this study were cultured in DMEM (10% FBS, 1×Antibiotic-Antimycotic) under 37 °C, 5% CO_2_.

#### Virus strains and cell infection

For SARS-CoV-2 infection, Huh7.5.1 cells were cultured in T-175 flasks (2×3×3 flasks, including 3 biological repeats, 3 flasks for each mock or infected ChIRP-MS), at a density of 5×10^6^ cells. The cells were briefly washed with DMEM at 16 h after seeding, and incubated with a clinical isolate of SARS-CoV-2/IPBCAMS-YL01/2020 for 1 h at a multiplicity of infection (MOI) of 0.05. Then the cells were supplemented with DMEM maintenance medium containing 1% FBS and cultured at 37°C, 5% CO_2_ for an additional 30 h. The cultured cells were washed twice with PBS and 4% formaldehyde (Pierce, 28908) was added for crosslinking at room temperature for 4 h. Live virus was inactivated by an additional 12 h incubation with 4% formaldehyde at 4 °C. The cells were then collected and washed with 0.125M Glycine at room temperature, followed by centrifugation at 1000g, 4 °C for 5 min and 3 washes with PBS. The cell pellets were used for ChIRP-MS experiments. Mock cells (no infection) were cultured and treated the same as the infected cells. All experiments involving live SARS-CoV-2 in this study were performed in a biosafety level III facility.

For ZIKV infection, 7×10^6^ Huh7 cells were cultured on a 15-cm dish for 20 h (2×3×3 plates, including 3 biological repeats, 3 plates for each mock or infected ChIRP-MS experiment), then infected with ZIKV (MR766, MOI 0.5). After 72 h, cells were collected using trypsin digestion and washed twice with cold PBS, followed by crosslinking in PBS containing 3% formaldehyde at room temperature for 30 min. Crosslinking was stopped by adding a 1/10 volume of room temperature 1.25M Glycine for 5 min. The cells were then washed with 3 times with PBS. After centrifugation, PBS was removed and the cell pellets were used for ChIRP-MS. Mock cells were cultured and treated like infected cells, but no virus was added.

EBOVΔVP30-GFP(*32*) (strain Zaire Mayinga) virions were generated using VP30 expressing Vero E6 cell lines, then infected Huh7.5.1 cells expressing VP30 (Huh7.5.1-VP30) at a MOI=0.1. For EBOV ChIRP-MS experiment, Huh7.5.1-VP30 cells were cultured on 15-cm plates (2×3×2 plates, including 2 biological repeats, 3 plates for each mock or infected ChIRP-MS) at a density of 6×10^6^ cells, then infected with EBOVΔVP30-GFP virus. After 72 h, infected and mock cells were collected and crosslinked as in the ZIKV experiment.

#### Identifying host proteins interacting with viral RNA by ChIRP-MS

ChIRP-MS was performed according to the previous report(*19*) with some modifications. Briefly, crosslinked cells were resuspended in 1 mL lysis buffer (∼100mg cells) containing 50cmM Tris–HCl, pHc7.0, 10cmM EDTA, 1% SDS, 0.1% sodium deoxycholate, 0.5% DDM and 0.1% NP-40) and sonicated. Probes (2 uL of 100 μM) tiling the whole genome of virus (Table S1) were used to capture vRNA-protein complexes. MyOne C1 beads (Invitrogen,65001) were blocked with 3% BSA and yeast tRNA (Solarbio, T8630), and 200 uL of C1 beads was used for each ChIRP experiment. The captured materials were firstly washed with lysis buffer for 2 times × 5min, then washed with ChIRP wash buffer (2×SSC, 0.5% SDS) for 4 times × 5min. The co-purified proteins were reverse crosslinked using 0.3M NaCl buffer (7.5cmM HEPES, pHc7.9, 12.5cmM D-biotin, 0.3M NaCl, 1.5cmM EDTA, 0.2% SDS, 5mM DTT, 0.075% sarkosyl and 0.02% sodium deoxycholate) at 70 °C with shaking for 1 h. Proteins were precipitated in 20% TCA, then the protein precipitate was resuspended in RIPA buffer (25mM Tris-HCl, pH 7.4, 1% NP-40, 0.5% DOC, 0.1% SDS). Protein samples were loaded onto an SDS-PAGE gel and visualized using silver staining. Protein bands were excised from the gel and de-stained following product instructions (Pierce Silver Stain for Mass Spectrometry, 24600). The excised gel bands were desalted, pH adjusted, and digested using Trypsin overnight. The digested peptides were extracted from the gel using 50% acetonitrile and 0.1% formic acid, and analyzed via mass spectrometry (Thermo Scientific Q Exactive). ChIRP-MS control experiments were performed on cells without viral infection (mock) by using the same set of probes.

To determine the viral RNA recovery rate for ChIRP-MS (%RNA retrieve), 10 μL of lysate (1%) was removed from 1 mL of sonicated cell lysate, and used as input sample. At the last wash, a 10 μL volume of wash buffer containing beads (1%) was removed as the eluate for RNA quantification. Samples (input and eluate) were suspended in 100 μL PK buffer (10 mM Tris–HCl pH 7.0, 100 mM NaCl, 0.5% SDS, 1 mM EDTA) containing 20 μg/mL proteinase K (Roche, 3115879001). The input and eluate samples were incubated at 50 °C for 45 min and then 95 °C for 10 min with mixing. Finally, 300 μL of TRIzol LS reagent was added to extract RNA following the manufacturer’s instructions. For both input and eluate, 2 μL of RNA was used for reverse transcription for cDNA synthesis using a PrimeScript RT reagent Kit (TAKARA, RR047A). For EBOV, sequence specific primers for the EBOV genome and *Gapdh* mRNA were used for reverse transcription. *q*PCR of viral RNA and *Gapdh* were performed using SYBR Green kits (TAKARA, RR420A) following the manufacturer’s instructions. The following primers were used for reverse transcription or *q*PCR: *Gapdh* Forward primer: ACACCCACTCCTCCACCTTTGAC; *Gapdh* Reverse primer: ACCCTGTTGCTGTAGCCAAATTC. ZIKV E Forward primer: CCGCTGCCCAACACAAGGTGAAG; ZIKV E Reverse primer: CCACTAACGTTCTTTTGCAGACAT. SARS-CoV-2 NP Forward primer: GGGGAACTTCTCCTGCTAGAAT; SARS-CoV-2 NP Reverse primer: CAGACATTTTGCTCTCAAGCTG. EBOV NP Forward primer: CCGTTCAACAGGGGATTGTTCG; EBOV NP Reverse primer: CTGCTGGCAGCAATTCCTCAAG. EBOV reverse transcription primer: CTCAGAAAATCTGGATGGCGCCGAGTCTC *Gapdh* reverse transcription prime: CTGAGTGTGGCAGGGACTCCCCAG

#### ChIRP-WB

For ChIRP-western blotting (ChIRP-WB), Huh7.5.1 (SARS-CoV-2), Huh7.5.1-VP30 (EBOV), or Huh7 (ZIKV) cells were cultured on ten 10-cm plates, infected with virus and crosslinked as above described. Then, ∼150 mg mock or infected cells were resuspended in 1 mL lysis buffer and sonicated. ChIRP experiments were performed as described above. 10 uL of cell lysate (per 1 mL, 1%) was removed as input. After washing, MyOne C1 beads were resuspended using 50 uL of 0.3M NaCl elution buffer (7.5cmM HEPES, pHc7.9, 12.5cmM D-biotin, 0.3M NaCl, 1.5cmM EDTA, 0.2% SDS, 5mM DTT, 0.075% sarkosyl and 0.02% sodium deoxycholate), boiled at 70 °C for 1 h and then at 95 °C for 30 min to elute proteins. 10 uL of the protein eluate and input samples were separated on SDS-PAGE gels and immunoblotted using the specific antibodies listed in Table S6.

#### ChIRP-MS data analysis

Raw mass spectrometry data were processed with Proteome Discover using the built-in search engine to search against the human proteome (Uniprot database). Viral proteins, including SARS-CoV-2 (NC_045512.2), ZIKV (NC_012532.1), and EBOV (NC_002549.1) proteins were downloaded from NCBI and added into the database manually. Proteomic data were filtered by applying a minimum Protein Score of 1.5. Viral RNA interacting proteins were scored with the MiST scoring algorithm(*57*) using default parameters. For SARS-CoV-2 and EBOV infected samples, proteins identified with a MiST score > 0.6 were considered to interact with viral RNA. For ZIKV infected samples, the cutoff MiST score was set as > 0.7. Every identified protein was assessed with the same parameters as in MiST, *i.e.*, “Abundance” to represent protein abundance in virus infected samples, defined as the mean of the bait-prey quantities (spectral counts divided by protein length) over all replicates for virus-infected samples; “Specificity” to measure the uniqueness of proteins identified in virus-infected samples compared with non-infected samples (mock), defined as the proportion of the abundance of virus infected samples compared to the abundance of non-infection samples; “Reproducibility” to evaluate the variance of each identified protein abundance among replicates, defined as the normalized entropy of the bait-prey quantities over all replicates for virus-infected samples. The proteins interacting with at least two viral RNAs were defined as common interacting proteins.

#### Differential protein expression analysis

To analyze differential protein expression of SARS-CoV-2 vRNA interacting proteins (n=143) across human tissues, we obtained protein abundance values in 29 human tissues from a proteomics dataset(*58*). Then the abundances of interacting proteins in each tissue were compared to their medians protein abundance among all 29 tissues. The significance *P* value of enrichment was calculated by Mann–Whitney U test and adjusted by FDR.

#### Protein domain analysis

To analyze whether there were possible enriched protein domains among the identified proteins, we annotated the domains of identified proteins using the Uniprot database. For each protein domain, we counted the number of identified proteins containing this domain. The enrichment (odds ratio) and significance *P* value of each domain among identified proteins, compared to all human proteins, was calculated by Fisher’s exact test.

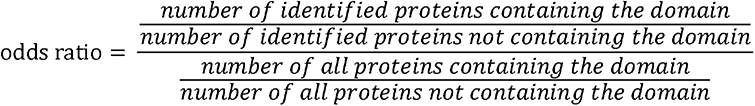

#### Protein complex analysis

To analyze whether there were possible enriched protein complexes among the identified proteins, we performed a protein complex enrichment analysis using the CORUM database(*59*). For each protein complex, we counted the number of identified proteins in this complex. The enrichment (odd ratio) and significance *P* value of these proteins in each complex, compared to all human proteins, was calculated by Fisher’s exact test.

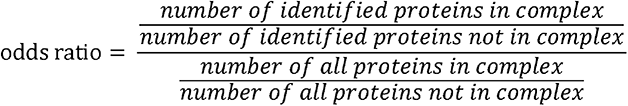

We searched the significantly enriched complexes (*P* value < 0.05) for each of the three viral RNA interactomes. Protein complexes present in at least two of the three virus interactomes were defined as common interacting complexes. Complexes present in only a single virus interactome were defined as virus-specific interacting complexes.

#### GO and KEGG enrichment analyses

We performed gene ontology (GO) and KEGG pathway enrichment analyses of the identified proteins using The Database for Annotation, Visualization and Integrated Discovery (DAVID) v6.8. The significance *P* values of GO terms and KEGG pathways were calculated by Fisher’s exact test. Top 10 enriched GO terms or KEGG pathways (*P* value < 0.05) were shown.

#### Interactome-informed drug discovery

SARS-CoV-2 host factors were assessed looking for drugs known to impact human proteins. Briefly, to identify drugs/compounds modulating 143 human interacting proteins, protein uniport IDs were searched against the IUPHAR/BPS Guide to Pharmacology, Drugbank, Drugcentral, and ChEMBL databases. Then we retrieved 5,309 compounds from these databases related to 56 host factors (Table S4). Specifically, 7 compounds related with 5 interacting proteins from the IUPHAR/BPS Guide to Pharmacology; 158 compounds related with 44 interacting proteins from the Drugbank; 19 compounds related with 6 interacting proteins from the Drugcentral and 5125 compounds predicted to be related with 35 interacting proteins from the ChEMBL (Table S4). Retrieved molecules were prioritized based on their FDA approval status and commercial availability. FDA approved drugs were prioritized for testing its antiviral activities; an exception is Silvestrol, which is an eIF4A specific inhibitor currently under investigation in a registered preclinical trial.

#### Antiviral drug screening

Based on the SARS-CoV-2 reference sequence (Wuhan-Hu-1, NC_045512), we cloned the SARS-CoV-2 genome as five fragments. These fragments were then assembled using *in vitro* ligation but replacing the viral N gene with GFP, to generate the SARS-CoV-2-GFPΔN genome. Genomic RNA of SARS-CoV-2-GFPΔN and mRNA of N gene were *in vitro* transcribed using an mMESSAGE mMACHINE T7 Transcription Kit (Thermo Fisher Scientific, AM1344). Synthesized viral RNA was electroporated into Vero E6-N cells that were stably expressing the SARS-CoV-2 N protein generated by lentivirus transduction, to produce the P_0_ virus. After 72 h, P_0_ virus was collected. Caco2 cells stably expressing the N protein (Caco2-N) were generated using lentivirus infection; this Caco2-N cell line can support the infection and replication of the SARS-CoV-2-GFPΔN virus.

To screen compounds with antiviral effects, Caco2-N cells were cultured on 96-well plates at a density of 1×10^4^ cells per well. After 24 h, cells were infected with SARS-CoV-2-GFPΔN virus at a MOI of 0.05 and analyte drugs were administered at the same time as the virus. After 72 h, FACS was performed to quantify GFP-expressing cells. Remdesivir was used as a positive antiviral control. The infection ratios resulting from treatment with various drug concentrations were normalized to the DMSO (0.2%) treated cells.

Compounds, known to target common interacting proteins and which showed antiviral effects against SARS-CoV-2-GFPΔN were also tested for EBOV and ZIKV infection. To assess the antiviral effects of the compounds on EBOV, Huh7.5.1-VP30 cells were cultured on 96 well plates with 1×10^4^ cells per well. After 20 h, drugs were added and cells were infected at the same time with EBOVΔVP30-GFP virus (strain Zaire Mayinga) at a MOI of 0.1. After 72 h, GFP-expressing cells were counted using FACS. Remdesivir was used as a positive antiviral control. The infection ratios with various drug concentrations were normalized to the DMSO (0.4%) treated cells.

For ZIKV, Huh7 cells were cultured on 96 well plates (Corning, 3603) with 1×10^4^ cells per well. To test the antiviral effects of drugs, Huh7 cells were infected with ZIKV (MR766) at a MOI of 0.5 and drugs were added at the same time. At 72 h, cells were fixed using 4% PFA, and blocked using blocking buffer (1×PBS, 5% FBS, 0.3% Triton X-100). The fixed cells were then incubated with Anti-Flavivirus Group Antigen Antibody (clone 4G2, MAB10216, 1:1000) in blocking buffer for 12 h at 4 °C. After washing 3 times with PBS, secondary antibody (Goat Anti-Mouse IgG H&L Alexa Fluor 488, ab150113) was diluted in blocking buffer (1:1000) containing DAPI (5 ug/mL) and was added and incubated for 2 h at room temperature. The ZIKV infection ratio was then quantified using an Opera Phenix High-content System. Total cell numbers were quantified based on DAPI staining of nuclei, and infected cells were quantified according the Alexa Fluor 488 staining in cytoplasm. The infection ratio was calculated as the number of Alexa Fluor 488 stained cells divided by the number of DAPI stained nuclei. The infection ratios with various drug concentrations were normalized to the DMSO (0.4%) treated cells.

For cell viability assays, Caco2-N, Huh7.5.1-VP30, and Huh7 were seeded on 96 well plates with 1×10^4^ cells per well, and treated using drugs with different concentrations. At 72 h, cell viability was measured using CellTiter-Glo Luminescent Cell Viability Assay kits (Promega, G7570). Cell viability values were normalized to DMSO treated cells (0.2% for Caco2-N, 0.4% for Huh7 and Huh7.5.1-VP30 cells).

Viral infection curves and cell viability curves were fitted using GraphPad Prism 8 (Nonlinear regression, Dose-response-Inhibition); the half maximal inhibitory concentration (IC50) and the half maximal cytotoxic concentration (CC50) values were calculated. The selective index (SI) was also calculated as the ratio of CC50/IC50.

## Supplementary Tables

**Table S1** Oligo probes used for ChIRP-MS

**Table S2** Viral RNA interacting proteins identified by ChIRP-MS

**Table S3** Enriched complexes of viral RNA interacting proteins

**Table S4** The information of drugs targeting SARS-CoV-2 RNA interacting human proteins from drug database

**Table S5** Information of the 21 selected drugs

**Table S6** Regents, cell line, virus and antibodies

## Supplementary Figures

**Fig. S1.**
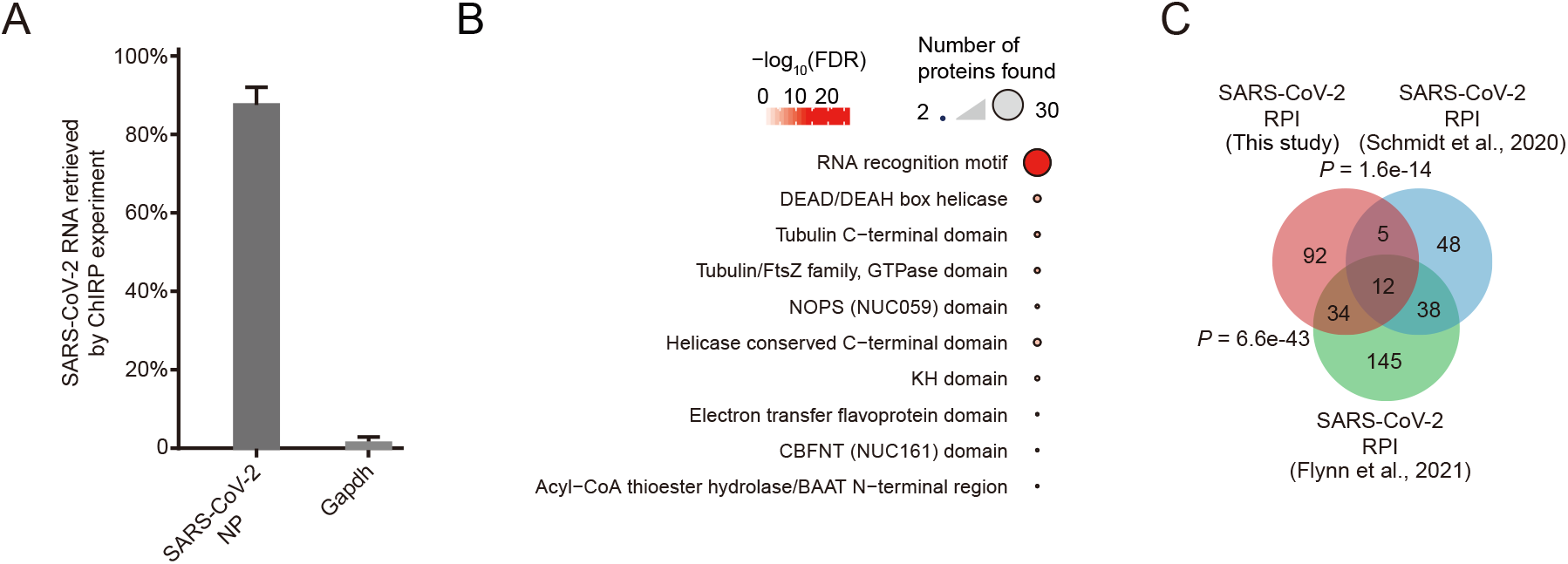
Identification of the human proteins interacting with SARS-CoV-2 vRNA in infected cells. **A,** SARS-CoV-2 vRNA recovery rates of ChIRP-MS experiments. *q*PCR of viral RNA (NP gene) was performed in ChIRP input and elution samples, *Gapdh* was used as a SARS-CoV-2 probe non-targeting control. Data are means ± SD, n= 3 biologically independent samples of ChIRP-MS. **B,** Protein domain enrichment analysis of the interacting proteins of SARS-CoV-2 vRNA. The top 10 enriched terms are shown. **C,** Comparison of SARS-CoV-2 vRNA interacting proteins identified in this study with protein groups from two different RNA pull-down method (*60, 61*) (Fisher’s exact test).

**Fig. S2.**
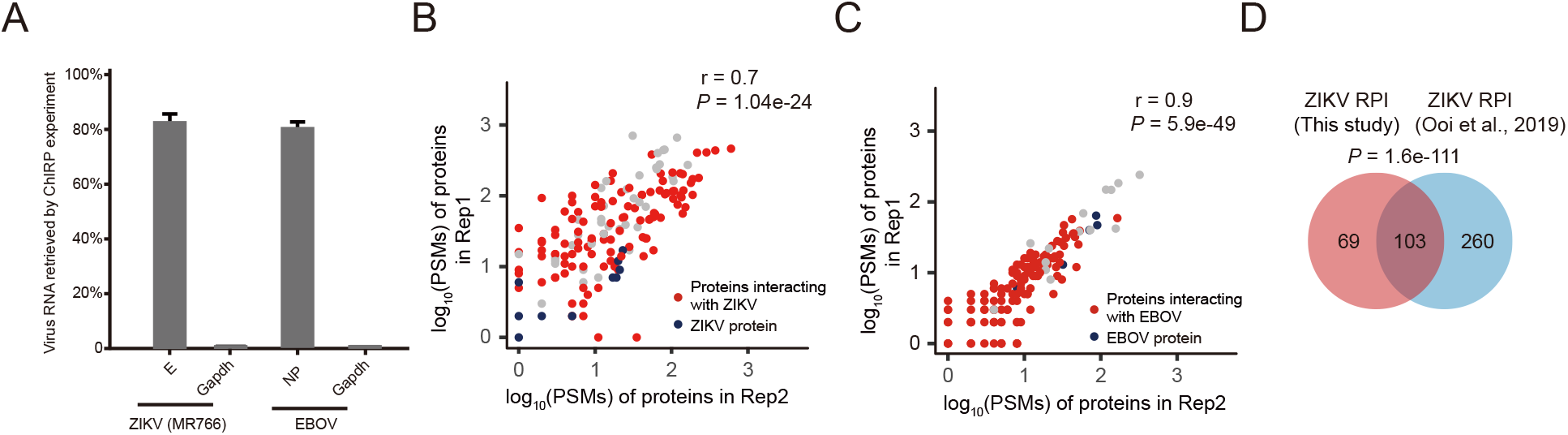
Identification of human proteins interacting with ZIKV and EBOV vRNAs in infected cells. **A,** Viral RNA retrieved by ChIRP-MS experiments. *q*PCR of viral RNA (ZIKV E protein gene and EBOV NP gene) was performed in ChIRP-MS input and elution samples, *Gapdh* mRNA was used as a non-targeting control for biotinylated oligo probes. Data are means ± SD, n=3 biologically independent samples for ZIKV, n=2 biologically independent samples for EBOV ChIRP-MS, respectively. **B, C,** Correlation of log_10_ (PSMs) for proteins identified in ZIKV infected Huh7 cells (**B**) and EBOV Δ VP30-GFP infected Huh7.5.1-VP30 cells (**C**), respectively. Data present Pearson correlation coefficient of log_10_ (PSMs) from biological replicates. The viral RNA interacting proteins are indicated using red dots in **B** and **C**; Viral proteins co-purified with viral RNA are in blue. **D,** ZIKV vRNA interacting proteins identified by different RNA pull-down methods. Human protein groups identified in the present study was compared with the protein groups identified by other studies(*17*).

**Fig. S3.**
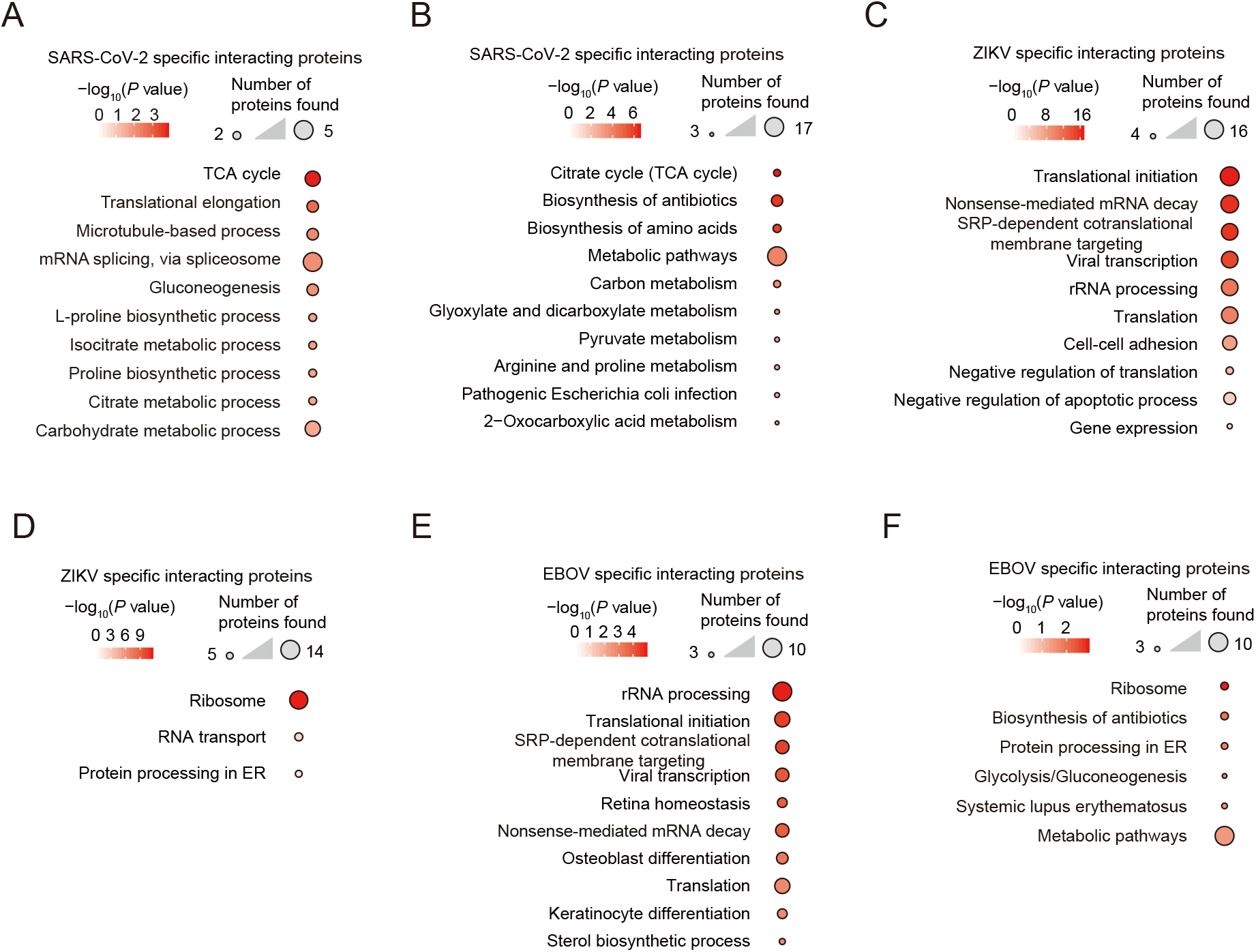
Functional analysis of the viral RNA-specific interacting proteins. **A, B,** Gene Ontology (**A**) and KEGG pathway (**B**) enrichment analysis of proteins specifically interacting with the SARS-CoV-2 vRNA. The top 10 enriched terms are shown. **C, D,** Gene Ontology (**C**) and KEGG pathway (**D**) enrichment analysis of proteins specifically interacting with the ZIKV vRNA. The top 10 enriched terms are shown. **E, F,** Gene Ontology (**E**) and KEGG pathway (**F**) enrichment analysis of proteins specifically interacting with the EBOV vRNA. The top 10 enriched terms are shown.

