## Supplemental Table S6 for "Analysis of viral RNA-host protein interactomes enables rapid antiviral drug discovery"

Table S6 Regents, cell line virus and antibodies

| REAGENT or RESOURCE | SOURCE | IDENTIFIER |
| --- | --- | --- |
| Antibodies | | |
| Cyclophilin A (PPIA) antibody | Proteintech | 10720-1-AP; RRID:AB_2237516 |
| Anti-IGF2BP1 (IMP1) (Human) pAb | MBL | RN007P; RRID:AB_1570640 |
| HSP90AB1 antibody | Proteintech | 11405-1-AP;  RRID:AB_2121207 |
| IDH2 antibody | Proteintech | 15932-1-AP;  RRID:AB_2264612 |
| Anti-hnRNPU | Proteintech | 14599-1-AP; RRID:AB_2248577 |
| SFPQ antibody | Proteintech | 15585-1-AP; RRID:AB_10697653 |
| Matrin 3 antibody | Abcam | ab151714; RRID:AB_2491618 |
| PTBP1 antibody | Proteintech | 55181-1-AP; RRID:AB_11182384 |
| Anti-Aly/REF | EMD Millipore | 03-120; RRID:AB_10807991 |
| NONO antibody | Proteintech | 11058-1-AP; RRID:AB_2152167 |
| ELAVL1 antibody | Abcam | ab200342;  RRID:AB_2784506 |
| PDIA6 antibody | Proteintech | 18233-1-AP; RRID:AB_10805765 |
| SND1 antibody | Abcam | ab65078;  RRID:AB_1566748 |
| RACK1 antibody | Santa Cruz | sc-17754; RRID:AB_2247471 |
| S100A9 antibody | Proteintech | 26992-1-AP; RRID:AB_2880716 |
| HIST1H3A antibody | Proteintech | 20532-1-AP;  RRID:AB_2811276 |
| TUBB antibody | Abcam | ab108342;  RRID:AB_10866289 |
| GAPDH (HRP Conjugated) | EASYBIO | BE0034-100;  N/A |
| SARS-CoV-2 spike protein antibody | GeneTex | GTX632604; RRID:AB_2864418 |
| Flavivirus 4G2 antibody | Millipore | MAB10216; RRID:AB_827205 |
| Bacterial and Virus Strains | | |
| SARS-CoV-2 | Gift | IPBCAMS-YL01/2020 |
| SARS-CoV-2-GFPΔN | This study | N/A |
| EBOVΔVP30-GFP (Mayinga) | Gift | From Ding lab |
| ZIKV MR766 | Dick et al., 1952 | N/A |
| Experimental Models: Cell Lines | | |
| Homo sapiens: Huh7 | Taguwa et al., 2015 | N/A;  RRID:CVCL_0336 |
| Homo sapiens: Huh7.5.1 | Gift | From Yang lab |
| Homo sapiens: Caco-2 | Gift | From Ding lab |
| Homo sapiens: A549 | ATCC | CCL-185;  RRID: CVCL_0023 |
| Cercopithecus aethiops: Vero | ATCC | CCL-81;  RRID: CVCL_0059 |
| Software and Algorithms | | |
| DAVID v6.8 | Dennis et al., 2003 | https://david.ncifcrf.gov/ |
| InterProScan | Jones et al., 2014 | https://www.ebi.ac.uk/interpro/ |
| Cytoscape | Shannon et al., 2003 | https://cytoscape.org/ |
| MiST | Jager et al., 2011 | https://github.com/everschueren/MiST |
